# A human model of the neurodevelopmental metabolic shift

**DOI:** 10.64898/2026.10.09.758091

**Authors:** Sydney N. Pettit, Ariella T. Kornfeld, Celeste Morales, Mollie K. Meffert

## Abstract

The neurodevelopmental metabolic shift transitions the metabolic pathways used by neurons to generate energy in order to critically support survival, growth, and functional maturation. To better understand metabolic regulation during human neuronal development and to facilitate the study of early neurodevelopmental disorders impacting this time window, establishing a combination of effective human model systems will be required. We report that a streamlined strategy using the genetic induction of neuronal differentiation by NGN2 expression recapitulates hallmarks of metabolic reprogramming associated with neurodevelopment. The approach is refined to increase the uniformity of differentiation, allowing reproducible high-throughput assessments. We validate appropriate readouts of metabolic function, including gene expression, cellular respiration, and mitochondrial morphology, which can be combined to present a cohesive assessment of metabolic status and reliance on oxidative phosphorylation relative to glycolysis. Collectively, the strategy presented will enhance the ability of investigators to gather foundational and disease-related insights into regulatory mechanisms of the metabolic shift critical to successful neuronal differentiation.

**Highlights:**

- Establishes a model of the neurodevelopmental metabolic shift using human i^3^Neurons
- Characterizes metabolic correlates across neuronal differentiation
- Demonstrates strategies to assess cellular metabolism and bioenergetics in hPSCs and neurons
- Provides a uniform system to probe metabolic deficiencies in neurological disorders

**Motivation:** Several approaches to investigate human neurodevelopmental metabolic reprogramming have been previously reported. However, prior strategies have largely relied on extrinsic factors to provide neurodevelopmental cues, which can impact the uniformity of differentiation as well as being costly, time-consuming, and often technically challenging. As impairments impacting cellular respiration impede proper neuronal development and metabolic deficiencies have been linked to neurodegenerative, neuropsychiatric, and neurodevelopmental disorders, there is a need for efficient and robust models to investigate these early developmental mechanisms. Here, we report a strategy, incorporating an established neuronal differentiation model using induction of stably integrated Neurogenin-2 (NGN2) for rapid generation of excitatory neurons (i^3^Neurons), which we demonstrate can synchronously recapitulate changes associated with the neurodevelopmental metabolic shift.

## INTRODUCTION

Glycolysis and mitochondrial oxidative phosphorylation (OXPHOS) are the two primary metabolic pathways used by cells to break down glucose for the production of Adenosine Triphosphate (ATP) and other metabolites needed for cellular survival, function, growth, and development.^1^ Glycolysis takes place in the cytoplasm and produces a net of two ATP, as well as NADH, carbon intermediates, and pyruvate. Pyruvate can subsequently either be reduced to lactate, which regenerates NAD+ to maintain glycolysis, or transported into the mitochondria where it is converted to acetyl-CoA and used by the tricarboxylic acid (TCA) cycle to produce the electron carriers, NADH and FADH2, which deliver their electrons to the electron transport chain (ETC) to power mitochondrial proton pumping and OXPHOS. Total cellular respiration generates 30 – 32 molecules of ATP per glucose, with OXPHOS contributing 28 – 30 of these ATP.^1^ Historically, pluripotent stem cells were thought to rely heavily upon aerobic glycolysis to produce ATP and necessary carbon intermediates, which serve as cellular building blocks in these dividing cells. However, recent studies reveal a more nuanced understanding of metabolism in stem cells, with several reports showing that mitochondrial OXPHOS is also high in naïve or early stem cells and that metabolic regulation then shifts across the spectrum of differentiation, from stem cells primed for differentiation, to committed progenitors, and finally to maturing neurons; however, this process is still not completely understood (**Figure 1A**).^2,3^

**FIGURE 1:**
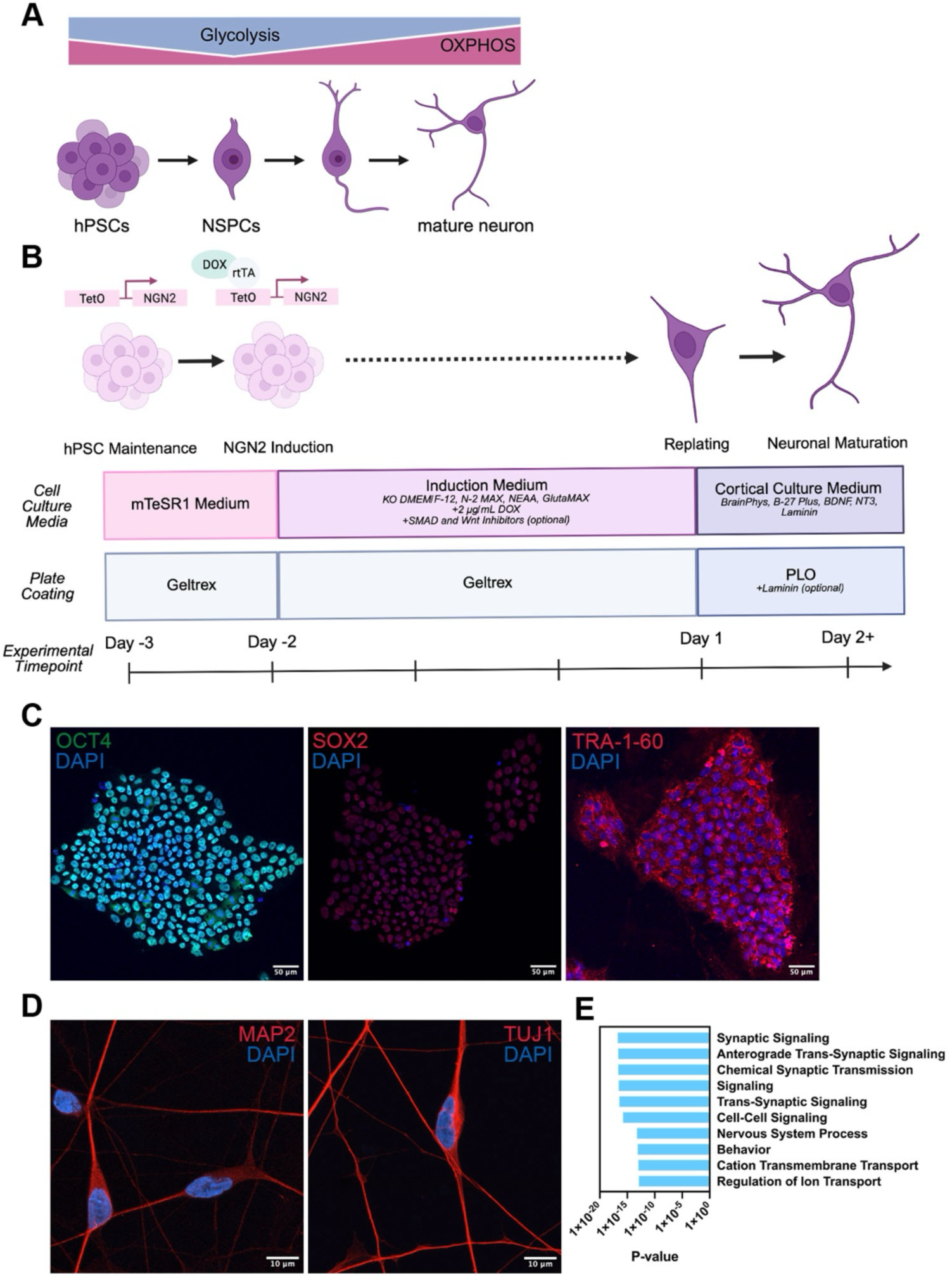
hPSC differentiation to i^3^Neurons. A) Schematic depicting the metabolic shift during neuronal differentiation from aerobic glycolysis in human pluripotent stem cells (hPSCs) and neural stem/progenitor cells (NSPCs) to predominance of mitochondrial oxidative phosphorylation (OXPHOS) in mature neurons. B) Schematic depicting i^3^Neuron differentiation workflow via doxycycline (DOX) inducible Neurogenin-2 (NGN2) and experimental time points. C) Representative immunocytochemistry (ICC) images showing hPSCs positive for pluripotent markers OCT4, SOX2, and TRA-1-60. D) Representative ICC images showing Day 7 i^3^Neurons positive for neuronal markers MAP2 and TUJ1. E) Gene ontology (GO) analysis of bulk RNA-seq data showing genes up-regulated during i^3^Neuron differentiation (Day 21); the top ten biological processes are ranked by p-value.

As neural stem/progenitor cells (NSPCs) mature into neurons, cells undergo a metabolic adjustment, shifting to prioritize OXPHOS for high efficiency of ATP production. Neurons require efficient ATP generation in order to match energy supply to rapid cellular demands for processes such as long-distance intracellular transport, synaptic transmission, and repolarization following action potential generation.^4–6^ Failure to produce sufficient ATP to meet the energy demands of the nervous system has been widely linked to bioenergetic failure in neurons, impairments in neural maturation, and cell death.^7,8^ While the metabolic shift from glycolysis to OXPHOS is tightly coupled with neuronal differentiation, the molecular mechanisms underlying this shift lack clarity. The neuronal metabolic shift has been shown to be heavily cell-intrinsic and studies have also indicated critical period differences between developing neurons in human and rodents.^9,10^ Collectively, prior work supports both the value and feasibility of defining human models of the metabolic shift in neurons which can be readily leveraged for greater mechanistic insights.

Neurodevelopment requires precise, and often localized, regulation of metabolic processes, including lipid, amino acid, and glucose metabolism, in order to meet the bioenergetic demands of maturing neurons and orchestrate proliferation, synaptogenesis, and neuronal activity.^11–15^ Disruption of these metabolic processes can result in a range of neurodevelopmental disorders including developmental delays, epilepsy, movement disorders, intellectual disabilities, and autism spectrum disorder (ASD).^16–23^ In neurons and other cell types, the metabolic shift can play an important role in the regulation of pluripotency and differentiation.^24^ Notably, metabolic dysfunction has been reported in patients and models of the neurodevelopmental disorder Fragile X syndrome (FXS), caused by reduced expression of fragile X messenger ribonucleoprotein (FMRP), and has also been suggested as an underlying molecular mechanism of the disorder.^25–30^ FMRP deficiency is characterized by aberrant synapse growth, neuronal hyperexcitability, and overabundant, immature dendritic spines.^31,32^ A mouse model exhibiting features of FXS showed altered synaptic mitochondrial morphology and respiration,^27,28^ a mitochondrial inner membrane leak linked to abnormal levels of the ATP synthase c-subunit, and an increase in the glycolytic enzymes hexokinase 2 (HK2) and lactate dehydrogenase (LDH).^28^ Trisomy of chromosome 21, resulting in Down syndrome, is another example of a neurodevelopmental disorder exhibiting widespread cellular bioenergetic deficiencies as well as specific links to aberrant mitochondrial metabolism. Down syndrome is the most frequent chromosomal abnormality resulting in intellectual disability, with many dosage-sensitive genes involved in mitochondrial function having been localized to chromosome 21 and exhibiting altered expression early in brain development.^33,34^

Metabolic reprogramming is integral to the proper development of mature neurons. Failure to complete the metabolic shift with appropriate timing can lead to an inadequate bioenergetic supply for developing neurons, resulting in cellular and network developmental defects. Validating an efficient human model with synchronous metabolic reprogramming in developing neurons offers the opportunity to reliably apply molecular and high throughput approaches to probe the mechanisms underlying this critical neurodevelopmental window, with the potential to reveal therapeutic targets for the treatment of neurological disorders associated with metabolic deficiencies. In this work, we use human embryonic stem cells (hESCs), a type of human pluripotent stem cell (hPSC), engineered with integrated inducible NGN2 as our model system.^35,36^ Leveraging the largely cell-intrinsic nature of the metabolic shift in neurons, we use a suite of established differentiation and metabolic markers to delineate the timing and optimize the approaches for investigations of the metabolic shift in a human neurodevelopmental model.

## RESULTS

### Establishing human neuron differentiation

To begin evaluating whether the neurodevelopmental metabolic shift could be recapitulated in the setting of developing human neurons, we first implemented and validated a robust strategy for the differentiation of neurons from hPSCs. This section describes the steps and critical parameters to establish neuron differentiation. A full benchtop protocol is described in a companion *STAR Protocols* publication.

Accurate large scale biochemical and molecular study of the metabolic shift requires that the developmental time course of neurons is as synchronous as possible. For this reason, a strategy incorporating NGN2 induction is implemented, which has been previously characterized for the efficient generation of i^3^Neurons (integrated, inducible, and isogenic human neurons).^35–38^ By using a stably integrated doxycycline-inducible NGN2 cassette (iNGN2), a scalable, homogenous neuronal maturation is achieved that is useful for high-throughput developmental and disease modeling. While non-integrating mRNA delivery can be employed, we recommend iNGN2 integration into a safe harbor locus (e.g. AAVS1 on human chromosome 19) using AAV-mediated iNGN2 delivery and stable line formation to increase the uniformity of differentiation.^37,39–41^ Our approach is based on existing protocols,^35–37^ but with modifications to promote synchronicity in the timing of differentiation and to limit the persistence of undifferentiated cells (**Figure 1B**). While NGN2 has been considered a master regulator of neurogenesis, which can rapidly generate neurons from hESCs or human induced pluripotent stem cells (iPSCs),^42^ a degree of neuron heterogeneity is nonetheless sometimes observed, and fate acquisition shows sensitivity to the dosage and duration of NGN2 expression, particularly when used in isolation.^43^ Coupling iNGN2 with the addition of patterning cues, in the form of small molecules inhibiting SMAD and WNT signaling, can improve the uniformity of excitatory neuron differentiation.^44^ The timeline of the strategy employed here is outlined in Figure 1B. While the approach we describe has been optimized using an established human embryonic stem cell line,^35,36^ it should be conceptually and technically applicable to other hPSC lines.

Using the WA01 (H1) hESC line engineered with stable incorporation of the iNGN2 cassette, we differentiated i^3^Neurons for seven days and conducted comparative testing by immunocytochemistry (ICC) of markers in both the stem cell line and i^3^Neurons to assess differentiation status; specifically, pluripotent markers OCT4, SOX2, and TRA-1-60 in hPSCs (**Figure 1C**) and neuronal markers MAP2 and TUJ1 in i^3^Neurons (**Figure 1D**) were compared by fluorescence microscopy, as these have been previously characterized as core markers for cell state characterization.^37,45–47^ The use of these markers in combination, rather than in isolation, further supports reliable determination of differentiation status. In addition, evaluation of residual undifferentiated cells or expression of pluripotency markers in i^3^Neurons is carried out by ICC for OCT4, SOX2, and TRA-1-60 (**Figure S1A**). Incorporation of nuclear staining (e.g. DAPI or Hoechst) in all ICC experiments enables identification of critical parameters, including in cells lacking pluripotent markers in hPSC cultures and cells post-differentiation which either exhibit pluripotent markers or fail to exhibit neuronal markers MAP2 and TUJ1.

Neuronal differentiation is associated with characteristic changes in gene expression profiles. We used bulk RNA-seq to compare transcriptomic signatures of hPSCs and i^3^Neurons by gene ontology (GO) analysis. Terms related to synaptic signaling were among the top enriched in i^3^Neurons (**Figure 1E**). Molecular functions, such as ion channel activity, and cellular components, such as neuron and synapse, were also identified as enriched terms in i^3^Neurons by GO analysis (**Figure S1B & C**), supporting efficient neuronal differentiation from hPSCs. Furthermore, neuronal genes, such as MAP2, SYN1, SLC17A7 (VGLUT1), and TUBB3, were significantly upregulated in Day 21 i^3^Neurons, whereas pluripotency genes, including HMGA2, MYCN, NANOG, POU5F1 (OCT4), and SOX2, were downregulated (**Figure S1D**). In particular, the pluripotency factor LIN28A can be noted as one of the most downregulated genes. The LIN28 RNA binding protein, well-characterized for promoting pluripotency, is also known for upregulating glycolysis in settings of cancer and for its roles in systemic glucose control and the modulation of insulin sensitivity.^48,49^ LIN28 mediates these functions in part through inhibiting the processing of let-7 family microRNA (miRNA) precursors to mature let-7 miRNAs, resulting in lowered let-7 miRNA levels, as well as by directly binding to mRNAs in some cases.^50^ Roles in both glucose metabolism and development mean that assessing the LIN28/let-7 developmental transition is, consequently, fundamental for establishing differentiation as well as the potential timing of a metabolic shift in developing neurons. LIN28 expression levels, including both LIN28A and LIN28B paralogs, are typically high in stem cells and decline during development to a low basal state in fully differentiated cells, while the let-7 miRNAs exhibit the opposite pattern (**Figure 2A**).^51^

**FIGURE 2:**
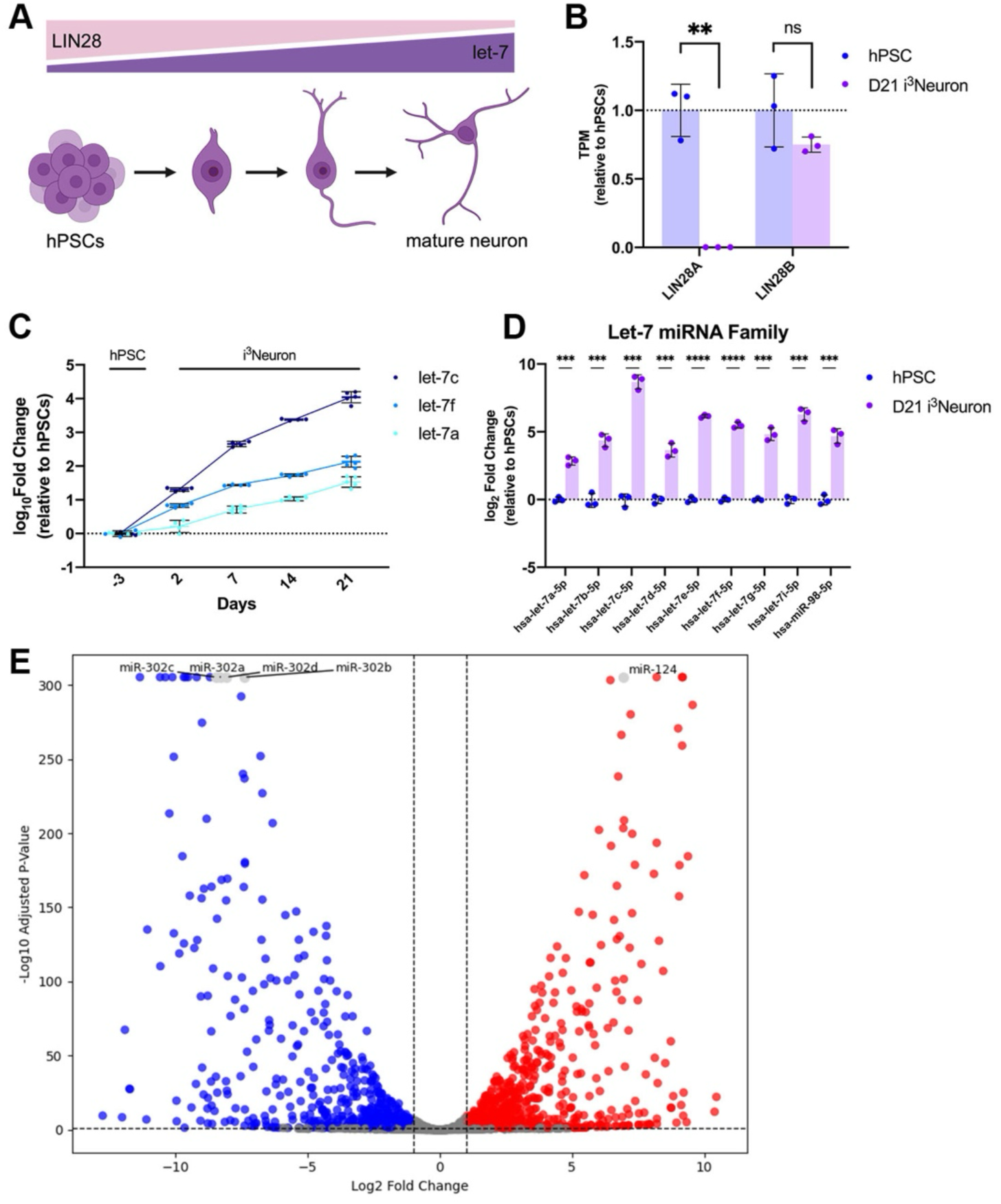
Changes in gene expression associated with neuronal differentiation in i^3^Neurons. A) Schematic depicting changes in the basal LIN28/let-7 axis during neuronal differentiation. B) Transcripts per million (TPM) values of LIN28A and LIN28B in Day 21 i^3^Neurons, relative to hPSCs [Unpaired t-tests with Holm-Sidak Multiple Comparisons; n= 3]. C) RT-qPCR data, normalized to small noncoding RNAs RNU48 and RNA5S using the ΔΔCt method (shown as log_10_ fold change), demonstrating expression of let-7 family members let-7c, let-7f, and let-7a in i^3^Neurons (Days 2–21) relative to hPSCs (Day -3) [n= 4–5]. D) Counts per million (CPM) values of let-7 miRNA family members in Day 21 i^3^Neurons relative to hPSCs (shown as log_2_ fold change) [Unpaired t-tests with Holm-Sidak Multiple Comparisons; n= 3]. E) Small RNA-seq data, following analysis with DESeq2, showing up-(red) and down-regulated (blue) miRNAs in Day 21 i^3^Neurons. Highlighted miRNAs are labeled and shown in light gray. Error bars represent mean ±SD, ****p< 0.0001, ***p< 0.001, **p< 0.01, ns= not significant.

Bulk RNA-seq showed that the LIN28A transcript was markedly reduced in Day 21 i^3^Neurons compared to the hPSCs (**Figure 2B**). Concordant with the downregulation of LIN28, we observed a marked elevation in members of the let-7 family (which share a consensus seed sequence) by RT-qPCR beginning at Day 2 and continuing to Day 21 in differentiating neurons, compared to hPSCs (Day -3) (**Figure 2C**). Small RNA sequencing (sRNA-seq) of hPSCs and i^3^Neurons at Day 21 validated the upregulation of multiple let-7 miRNA family members (**Figure 2D**). sRNA-seq also demonstrated robust changes in miRNA expression, including upregulation of the neuron-specific miRNA, miR-124,^52^ which is the most abundant miRNA in the brain and expected to be enriched in successfully differentiated i^3^Neurons (**Figure 2E**). In addition, sRNA-seq data showed that members of the miR-302 family, embryonic stem cell-specific miRNAs,^53^ were enriched as expected in the hPSCs (**Figure 2E**). Collectively, evaluation of these data supported successful neuronal differentiation by the strategy described. These data also suggested that we might further evaluate this setting as a possible model system for study of the metabolic shift in human neuronal development.

### Assessing metabolic gene reprogramming in human neurons

Previous studies have linked let-7 and miR-124 miRNAs to the regulation of metabolism.^48,49,54,55^ However, while these miRNAs are known to promote neurogenesis and neuronal differentiation,^51,52^ their potential roles on metabolic reprogramming during neurodevelopment has not been well characterized. Consistent with the upregulation of let-7 miRNAs during neuronal differentiation, evaluation of bulk RNA-seq indicated that several known metabolic gene targets of let-7 miRNAs, including neuroblastoma-derived myelocytomatosis (N-MYC)^56,57^ and hexokinase 2 (HK2),^58^ were significantly downregulated in i^3^Neurons at Day 21 of differentiation, relative to hPSCs (**Figure 3A**). N-MYC is a growth-promoting transcription factor known for its multifaceted functions in enhancing glucose uptake and boosting glycolysis,^59^ while HK2 is the initial and rate-limiting enzyme in glycolysis.^1^ These data were in-line with metabolic reprogramming occurring by Day 21 of neuronal differentiation in this system. Peroxisome proliferator-activated receptor gamma coactivator 1-alpha (PGC-1ɑ), encoded by the *PPARGC1A* gene, is a transcriptional co-activator that promotes mitochondrial biogenesis and is known for promoting oxidative metabolism at the expense of glycolysis.^15,60^ PGC-1ɑ can maintain the transcription of multiple nuclear-encoded mitochondrial genes including the mitochondrial transcription factor A (TFAM), which then works inside the mitochondria to initiate mitochondrial transcription.^11,12^ We noted that the PGC-1ɑ transcript was also significantly upregulated in Day 21 i^3^Neurons (**Figure 3B**). Further, estrogen-related receptor gamma (ERRγ), a nuclear receptor and transcription factor that interacts with PGC-1ɑ to promote mitochondrial transcription,^61^ is also significantly increased in i^3^Neurons, while other factors (nuclear respiratory factor 1 (NRF1) and mitochondrial RNA polymerase (POLRMT)) exhibit no changes in expression. Surprisingly, TFAM appears to be slightly downregulated (**Figure 3B**). However, as TFAM is a critical and therefore highly regulated protein,^62^ this apparent discrepancy could reflect regulation at the post-transcriptional level and may not be representative of changes in protein expression.

**FIGURE 3:**
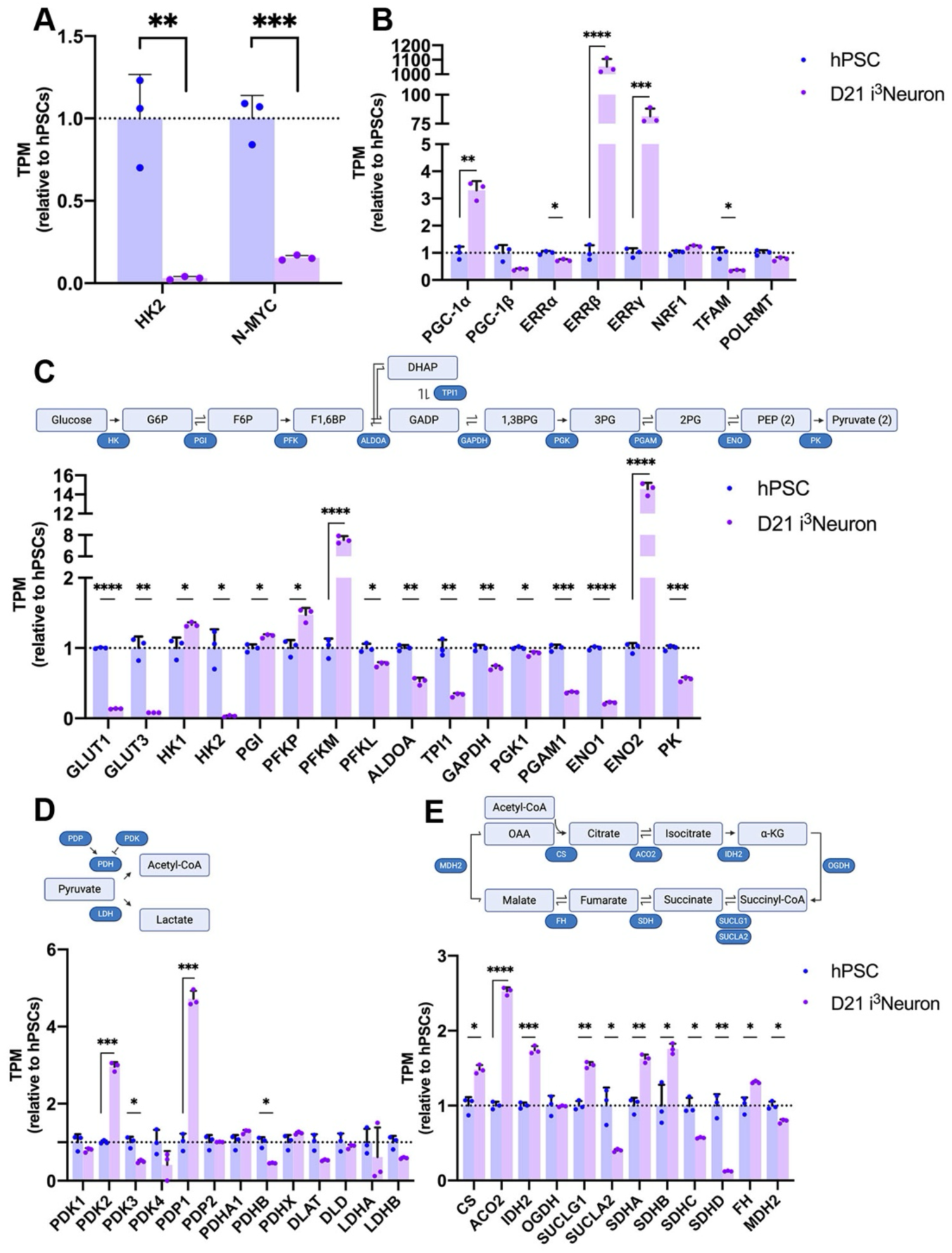
Regulation of metabolic gene expression associated with metabolic reprogramming in i^3^Neurons. A–E) Transcripts per million (TPM) values of A) HK2 and N-MYC; B) factors involved in the transcription of metabolic and mitochondrial genes; C) glycolytic genes; D) genes involved in pyruvate metabolism; E) tricarboxylic acid (TCA) cycle genes in Day 21 i^3^Neurons, relative to hPSCs [Unpaired t-tests with Holm-Sidak Multiple Comparisons; n= 3]. Error bars represent mean ±SD, ****p< 0.0001, ***p< 0.001, **p< 0.01, *p< 0.05.

Assessment of genes involved in critical steps of glycolysis (**Figure 3C**), pyruvate metabolism (**Figure 3D**), and the TCA cycle (**Figure 3E**) provide an additional means to evaluate the occurrence of a metabolic shift in i^3^Neurons. Many genes involved in glycolysis demonstrate a decrease in expression, including a loss in glucose transporters GLUT1 and GLUT3 (**Figure 3C**). A significant increase was observed for enolase 2 (ENO2), which is highly expressed in mature neurons,^63^ hexokinase 1 (HK1), and phosphofructokinase (PFKM), which is consistent with previous reports (**Figure 3C**).^15^ Furthermore, we note that genes involved in pyruvate metabolism also exhibit changes in expression across differentiation as expected (**Figure 3D**). Subunits of the pyruvate dehydrogenase complex (PDC), PDHA1, PDHX, DLAT, and DLD, which convert pyruvate to acetyl-CoA, show no significant changes in expression, with the exception of PDHB, which slightly decreases. Pyruvate dehydrogenase kinases (PDKs) inhibit the PDC ^64^ and appear to decrease, except for PDK2, which significantly increases in i^3^Neurons in concordance with other models.^15^ Furthermore, pyruvate dehydrogenase phosphatase 1 (PDP1), which activates the PDC,^64^ significantly increases. Lactate dehydrogenases LDHA and LDHB also appear to be slightly, although not significantly, decreased (**Figure 3D**). Finally, we note that the TCA cycle, which provides many important biosynthetic intermediates,^1^ also shows evidence of changes in gene expression consistent with metabolic reprogramming (**Figure 3E**). A majority of related genes appear to significantly increase in expression, besides succinyl-CoA ligase 2 (SUCLA2), succinate dehydrogenase complex subunit C and D (SDHC and SDHD), and malate dehydrogenase 2 (MDH2) (**Figure 3E**). These changes in metabolic gene expression, normalized to TPM, were broadly consistent with DESeq2 analysis (**Table S1**) and further support the occurrence of metabolic reprogramming in this model as well as provide information on the potential timing of the metabolic shift.

### Bioenergetic assessment of metabolic reprogramming

We next used a combination of bioenergetic assays to directly test the potential metabolic transition from predominant reliance on oxygen-independent glycolysis in hPSCs to increased utilization of mitochondrial OXPHOS during neuronal differentiation. The metabolic processing of glucose into pyruvate by cytoplasmic glycolysis subsequently leads to the enzymatic conversion of pyruvate into lactate in proliferating cells (**Figure 4A**),^65^ allowing the measurement of lactate levels to provide comparative assessments of relative glycolytic activity. Accordingly, we used a bioluminescence assay (Lactate-Glo) to monitor secreted L-lactate levels in hPSCs and across the neuronal differentiation timeline, with normalization to cell numbers using a live-cell nuclear stain (CyQUANT). Lactate secretion was strikingly decreased in i^3^Neurons compared to hPSCs, which began shortly after iNGN2 induction (Day 2), and was maintained until Day 21, the last timepoint measured (**Figure 4B**), consistent with a shift away from aerobic glycolysis and towards OXPHOS in i^3^Neurons. Measurement of the extracellular acidification rate (ECAR) by Seahorse XF Analyzer, provides an orthogonal perspective on the rate of glycolysis, as cells pump out both protons and lactic acid at elevated rates when relying on glycolysis for energy production. This data also supports a robust decrease in the basal ECAR of Day 21 i^3^Neurons compared to hPSCs (**Figure 4C**).

**FIGURE 4:**
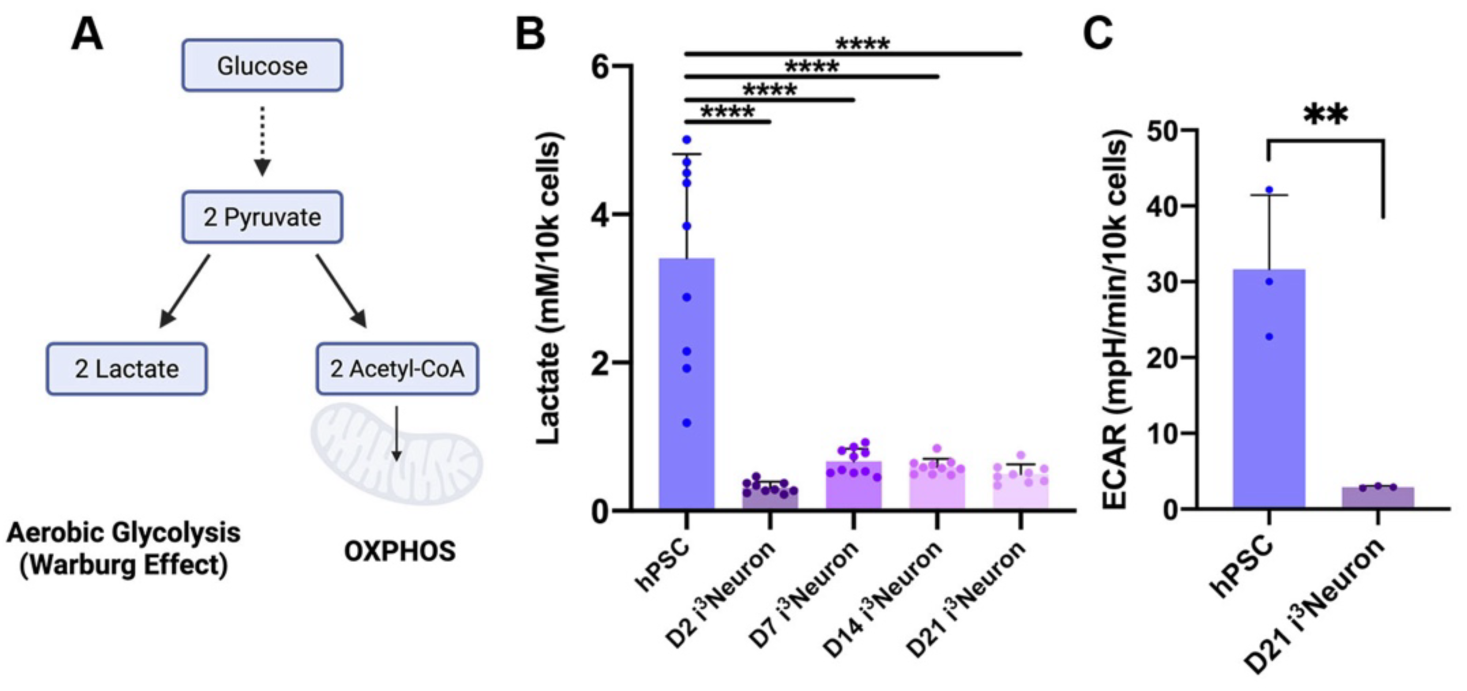
Functional readouts of aerobic glycolysis across differentiation of i^3^Neurons. A) Schematic depicting the metabolism of glucose to lactate during aerobic glycolysis (Warburg Effect) or to acetyl-CoA during OXPHOS. B) Lactate secretion (mM), measured using a Promega Lactate-Glo assay and normalized by cell number [One-way ANOVA; Dunnett’s Multiple Comparisons; n= 9–10]. C) Basal extracellular acidification rate (ECAR), measured by Seahorse XF Analyzer and normalized by cell number, in hPSCs and Day 21 i^3^Neurons. Each dot represents one experimental replicate, with 30–60 biological replicates per experiment [Unpaired t-test]. Error bars represent mean ±SD, ****p< 0.0001, **p< 0.01.

The analysis of respiratory parameters by Seahorse XF Analyzer also permits assessment of the relative oxygen consumption rates (OCR, pmol/min) and the response to standard injected inhibitors and uncouplers, which target specific components of the mitochondrial electron transport chain (ETC). To probe mitochondrial contributions to cellular respiration, we conducted a Seahorse XF Mitochondrial Stress Test using oligomycin to target ATP synthase, FCCP as a proton gradient uncoupler, and rotenone/antimycin A (Rot/AA) to eliminate mitochondrial contributions to OCR in hPSCs and Day 21 i^3^Neurons (**Figure S2A**). Data from this analysis can be used to quantify basal respiration, maximal respiratory capacity, spare respiratory capacity, ATP-linked respiration, and non-mitochondrial respiration. Interestingly, hPSCs exhibit significantly higher OCR than i^3^Neurons (**Figure S2B & C**). Accordingly, we observe a decrease in maximum respiration, spare respiratory capacity, and ATP-linked respiration in Day 21 i^3^Neurons (**Figure S2C – F**). This finding prompted further investigation into the concentration of intracellular ATP using a bioluminescent assay (Cell Titer-Glo). This data confirmed Seahorse Mitochondrial Stress Test results and indicated a higher total production of ATP per cell in hPSCs relative to Day 21 i^3^Neurons (**Figure S2G**).

We did observe significant changes, however, in the fraction of ATP production linked to OXPHOS or glycolysis in response to differentiation. In the i^3^Neurons, about 69% of the total ATP-linked production arose from mitochondrial (OXPHOS) contributions, compared to 54% in hPSCs (**Figure 5A**). This finding is consistent with reported changes in mitochondrial networks supporting neuronal bioenergetic requirements.^2^ To further investigate this aspect, we stained mitochondria, using MitoBrilliant, in hPSCs and i^3^Neurons and conducted super-resolution confocal imaging to compare mitochondria between differentiation states (**Figure 5B**). Quantification of mitochondria in cell bodies revealed that the mitochondria density (fraction of mitochondria area relative to cytoplasm area) was significantly greater in i^3^Neurons compared to hPSCs (**Figure 5C**). Taken together, our data indicate a shift towards a predominance of OXPHOS for energy production, and an expansion of the mitochondrial network. These results are consistent with the known reliance of neurons on high-efficiency mitochondrial mechanisms for energy production, which can also be locally adapted to fuel processes such as ion pumping for electrical signaling, membrane repolarization, and synaptic transmission.

**FIGURE 5:**
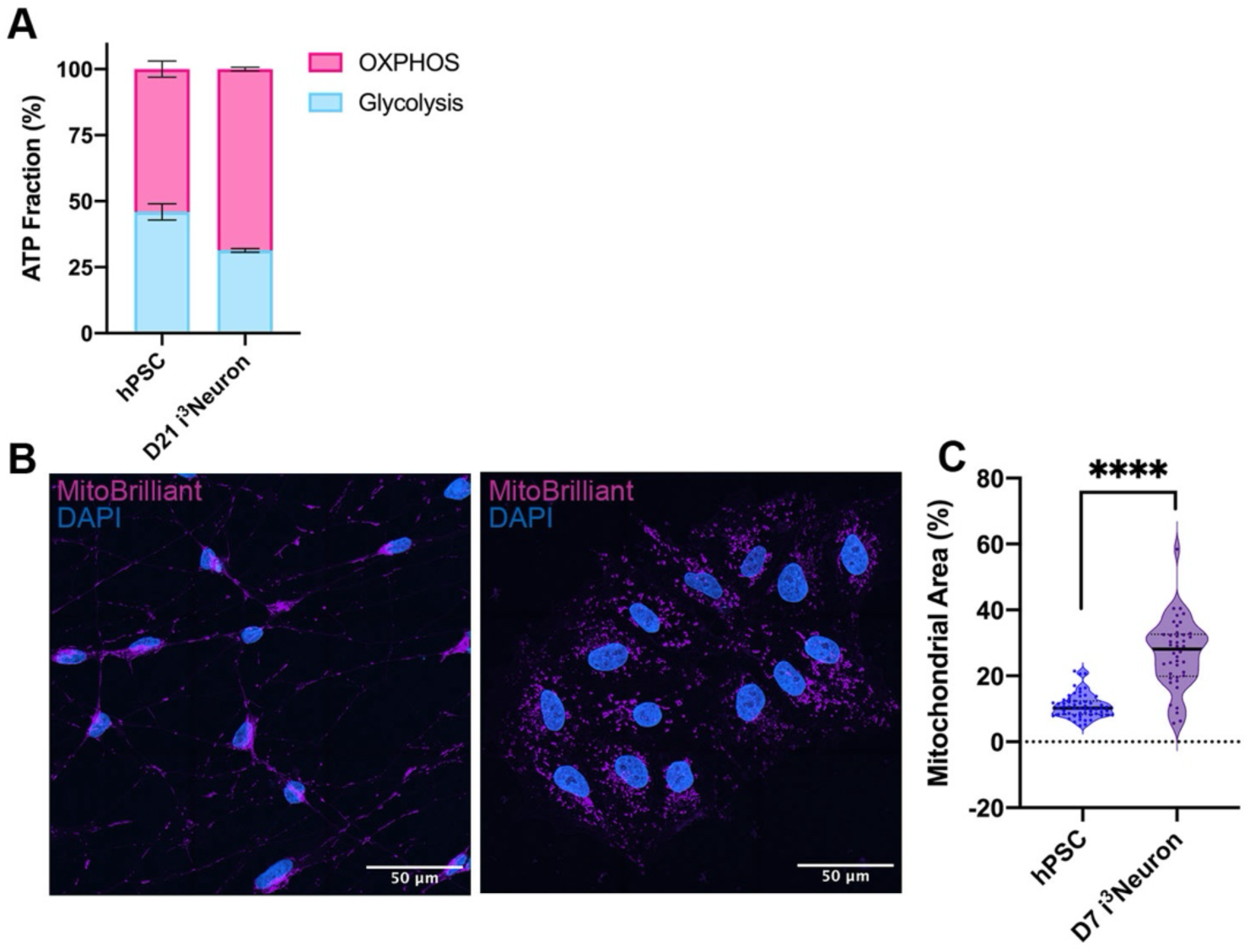
Shifts in cellular respiratory metabolism and mitochondrial morphology across differentiation of i^3^Neurons. A) Fraction of ATP production linked to glycolysis and mitochondrial OXPHOS in hPSCs (46% and 54% respectively) and Day 21 i^3^Neurons (31% and 69% respectively), determined using Seahorse XF Analyzer. Data includes three experimental replicates, with 30–60 biological replicates per experiment. B) Representative images showing mitochondrial morphology and density in Day 7 i^3^Neurons (*left*) and hPSCs (*right*) using MitoBrilliant 646 dye. C) Quantification of percentage of mitochondrial area per cell body area in hPSCs and i^3^Neurons (*as in* Figure 5B). Solid lines represent the median; dotted lines represent the first and third quartiles. Each dot represents one cell [Unpaired t-test; n= 42–59]. ****p< 0.0001.

### Identification of inadequate metabolic reprogramming

In this section we discuss aspects which, if not controlled, could contribute to failure of this model system to reliably recapitulate features of the neurodevelopmental metabolic shift. For purposes such as drug screening or the study of genetic disorders of the nervous system, uniformity in the timing of the developmental metabolic shift is critical for robust comparisons in high throughput assays. We find that the two primary sources of non-uniformity in development are: *i)* spontaneous differentiation in hPSC cultures, and *ii)* the persistence of populations of undifferentiated cells, or “de-differentiation,” in i^3^Neuron cultures.

Spontaneous differentiation in hPSC cultures can be detected by routine visual inspection under brightfield microscopy (example in **Figure S3A & B**), and further validated using staining for pluripotent markers, as described (**Figure 1C**). If observed, the following measures should be taken prior to proceeding with neuronal differentiation. If possible, the spontaneously differentiated hPSC culture should be entirely discarded, and a new vial with minimal differentiation thawed and used for experiments and i^3^Neuron differentiation. The presence of low-level spontaneous differentiation (≤ 10%, example in **Figure S3A**) is not uncommon and will not significantly interfere with neuronal differentiation, as we note that spontaneously differentiated hPSCs are typically lost over time. Moderate differentiation can often be addressed by gently passaging hPSCs and seeding them sparsely, since differentiated cells more strongly adhere to the plate surface, and rapidly dividing hPSCs will outcompete differentiated cells. Severe spontaneous differentiation (example in **Figure S3B**) should be avoided and if observed, cells discarded. If frozen hPSC vials are limited, manually picking undifferentiated hPSC colonies to rescue the culture can be performed; however, this is a severe selection and not recommended due to potential downstream consequences.

The persistence of undifferentiated populations in i^3^Neuron cultures can also be detected by routine visual inspection under brightfield microscopy (example in **Figure S3C**), and can be validated using staining for neuronal markers, as described (**Figure 1D**). We note that overly confluent hPSC cultures during iNGN2 induction often result in poor neuronal differentiation efficiency, so hPSCs should be only 70 – 80% confluent and appear healthy, with limited spontaneous differentiation, before proceeding with neuronal differentiation. We also note that excessive mechanical stress, such as pipetting, can result in i^3^Neuron death, clumping, detachment from the plate surface, and poor neuronal differentiation. Attention to reducing mechanical stress is particularly important for assays in 96-well plates, such as the Seahorse XF Analyzer and bioluminescent assays, which require multiple dilutions, seeding, and media changes with a multichannel pipet. The inclusion of small molecule inhibitors is strongly recommended for these applications, as they promote a more uniform neuronal differentiation, compared to doxycycline alone.^44^ Furthermore, coating plates with laminin, in addition to the usage of poly-L-ornithine, is recommended to strengthen adherence and reduce clumping of i^3^Neurons.^37^

## DISCUSSION

Early development in the human nervous system features unique progenitor cell dynamics, compared to other mammalian systems, and inherent differences in human-specific neuronal gene expression. In addition, cellular respiration is critical for proper neuronal development and function, although the underlying mechanisms linking these two processes are still not well understood. These considerations highlight the imperative for diverse human models of neuronal development to enable understanding of critical regulatory steps and to advance the discovery of mechanisms underlying dysregulation in neurodevelopmental disorders. Here, we define a human model using hPSC differentiation to neurons through iNGN2, which recapitulates changes associated with the neurodevelopmental metabolic shift. Collectively, this work highlights the i^3^Neuron system as a valuable model for studying metabolic reprogramming during the development of human neurons.

We demonstrate cell-intrinsic changes associated with neurodevelopment and the metabolic shift across neuronal differentiation in the i^3^Neuron model. Maturing neurons exhibit an increase in the fraction of ATP generation arising from OXPHOS compared to hPSCs (**Figure 5A**). We also observed that hPSCs exhibit high rates of OXPHOS and aerobic glycolysis (**Figure S2B–F** and **Figure 4**). This is consistent with previous reports which suggest that although NSPCs exhibit a lower requirement for OXPHOS relative to neurons,^66^ hPSCs rely on both glycolysis and mitochondrial respiration for high energy production.^2,3^

Attention to hPSC passage number, media composition, and the careful design of experiments is required to minimize variability, ensure reproducibility, and support robust interpretation of metabolic data in this system. For example, in embryonal carcinoma stem-like cells, metabolism has been linked to passage number, suggesting that variability in hPSC bioenergetics could be impacted by altered genomic integrity.^67^ A shift in metabolic rate has also been linked to glucose concentration of cell culture media.^68^ Our i^3^Neuron medium (BrainPhys) contains 2.5 mM glucose, which is comparable to physiological levels; however, the glucose concentration of many neuronal media formulations, such as Neurobasal, is much higher (25 mM), which could alter glucose metabolism and mitochondrial respiration compared to in vivo settings. Additionally, normalization to cell number is particularly important in assays comparing different stages of differentiation, and normalization strategies must be incorporated when quantifying differences in respiration. Due to the larger size and protein content of neurons relative to hPSCs and NSPCs, normalization to protein is not ideal. Normalization to DNA content using nuclear stains (e.g. CyQUANT) is an effective approach but requires awareness of differences in plate materials and dimensions, temperature variation, background fluorescence, well-to-well crosstalk, and the construction of standard curves to calculate cell number.

i^3^Neurons offer advantages as a model system, compared to human immortalized cell lines, neuroblastoma lines, or animal models, due to their ease of experimental manipulation, synchronized development, lack of transformation, relative cellular homogeneity, and ability to recapitulate aspects of normal human neuronal physiology and disorders. Computational modeling utilizing published data has been implemented in order to study human neuronal metabolism in genome-scale metabolic models.^69,70^ While computational approaches can capture disease-associated metabolic differences in human neurons they may be less useful for modeling chronologically progressive processes, such as the neurodevelopmental metabolic shift, which benefits from direct temporal comparisons. Studies of neuronal development between mouse and human nervous systems indicate inherent differences in gene expression and tempo, underscoring the value of evaluating healthy and disease processes in a human system.^3,9^ Data on the timing of the metabolic shift in the intact human nervous system is limited and relies primarily upon advanced neuroimaging techniques (such as in vivo Magnetic Resonance Spectroscopy) and ex vivo metabolic profiling from fetal tissue. Such studies lack cellular and temporal resolution, but have indicated dramatic metabolic changes between birth and three months largely focusing on whole brain and brain region changes in glucose metabolism.^71,72^ Experimental models including both 2D and organoid approaches have also been developed to model early human neocortical development from neuroepithelial stem cells, iPSCs, or hESCs.^15,73–75^ Typically, neuronal differentiation in these systems has been induced by a program of growth factors and small molecules, with comparison of differentiated neurons to neural progenitor cells rather than embryonic stem cells.^15^ While the timing and synchrony of development are not directly comparable between earlier studies and the later-developed i^3^Neuron system,^35–37,44^ parallels in markers of the metabolic shift towards a dominance of OXPHOS during neurodevelopment are observed. Finally, the approach described here may prove particularly valuable for studying largely cell-autonomous mechanisms in neurodevelopment, such as metabolic reprogramming, but could be coupled with additional relevant cell types in future studies.

### Limitations of the study

We demonstrate a shift in the fraction of total ATP production linked to mitochondrial OXPHOS and a decreased fractional reliance on glycolysis and total respiratory activity in Day 21 i^3^Neurons relative to hPSCs. Comparison to metabolic assays conducted at an earlier timepoint during i^3^Neuron differentiation, using a stage resembling neural stem/progenitor cells (NSPCs) rather than hPSCs, might reveal higher relative respiratory measurements in i^3^Neurons.

Previous studies indicate that hPSC-derived neurons may be similar in some aspects to neurons at prenatal stages rather than neurons at adult stages.^76^ Although substantial metabolic reprogramming appears to occur early on in neuronal differentiation according to our data and other reports,^15^ continued culture of i^3^Neurons could be done to investigate metabolic processes later in development.

In this study, we utilize a hESC-derived i^3^Neuron model system; although it seems likely that metabolic reprogramming would also be observed in iPSC-derived i^3^Neurons, we do not test this here. Since iPSCs have been shown to be capable of retaining gene expression and epigenetic signatures from their tissue of derivation and developmental stage,^77^ there may be differences in metabolic gene expression which impact cellular metabolism.

Due to the lack of other cell types and features found in the intact brain, this model may miss molecular interactions and processes that occur in vivo, which could be complemented by orthogonal follow-up experiments using animal models. To better understand metabolism during human brain neurodevelopment, assessing data from a combination of diverse human model systems will be necessary.

## Resource availability

## Lead contact

Further information and requests for resources and reagents should be directed to and will be fulfilled by the lead contact, Mollie K. Meffert.

## Materials availability

This study did not generate new unique reagents.

## Data and code availability

Raw FASTQ files for RNA-seq data have been deposited at GEO and are publicly available as of the date of publication. Accession numbers are listed in the key resources table.

This paper does not report original code. RNA-seq data was analyzed using existing code, which is listed in the key resources table.

Any additional information required to reanalyze the data reported in this paper is available from the lead contact upon request.

## Acknowledgments

We thank members of the Meffert laboratory for helpful discussions and critical feedback on the manuscript. We thank L. Barrett and members of the Barrett lab for generously sharing cell lines and culture expertise. We thank S. Claypool and the Claypool laboratory for sharing expertise in seahorse metabolic assays, and the J. Lynch lab for sharing expertise in fluorescence and luminescence detection strategies. Graphics generated using BioRender.

## Author contributions

S.N.P. and M.K.M. participated in project conceptualization and experimental design; S.N.P., A.T.K., and C.M. conducted experiments, replicated results, and prepared figures; S.N.P. and A.T.K. performed computational analysis; S.N.P., A.T.K., M.K.M. participated in manuscript drafting and revision; S.N.P. and M.K.M. participated in supervision.

## STAR★Methods

## Key resources table

| REAGENT or RESOURCE | SOURCE | IDENTIFIER |
| --- | --- | --- |
| Antibodies |  |  |
| Goat Polyclonal Anti-OCT3/4 | R&D Systems | Cat #AF1759; RRID: AB_354975 |
| Mouse Monoclonal Anti-SOX2 (245610) | R&D Systems | Cat# MAB2018; RRID: AB_358009 |
| Mouse Monoclonal Anti-TRA-1-60 | Santa Cruz Biotechnology | Cat# sc-21705; RRID: AB_628385 |
| Rabbit Polyclonal Anti-MAP2 | Sigma-Aldrich | Cat# AB5622; RRID: AB_91939 |
| Mouse Monoclonal Anti-Tubulin $\beta$ 3 (TUBB3) | BioLegend | Cat# 801213; RRID: AB_2313773 |
| Donkey anti-Goat IgG (H+L) Cross-Adsorbed Secondary Antibody, Alexa Fluor 488 | Invitrogen | Cat# A-11055; RRID: AB_2534102 |
| Goat anti-Mouse IgG (H+L) Highly Cross-Adsorbed Secondary Antibody, Alexa Fluor 568 | Invitrogen | Cat# A-11031; RRID: AB_144696 |
| Donkey anti-Rabbit IgG (H+L) Highly Cross-Absorbed Secondary Antibody, Alexa Fluor 568 | Invitrogen | Cat# A10042; RRID: AB_2534017 |
| Chemicals, peptides, and recombinant proteins |  |  |
| Geltrex™ LDEV-Free, hESC-Qualified, Reduced Growth Factor Basement Membrane Matrix | Gibco | Cat# A1413302 |
| DMEM/F12 | Gibco | Cat# 11320033 |
| mTeSR™1 Complete Kit | Stem Cell Technologies | Cat# 85850 |
| Normocin | InvivoGen | Cat# ant-nr-1 |
| Geneticin™ Selective Antibiotic (G418 Sulfate) | Gibco | Cat# 10131035 |
| Y-27632 dihydrochloride (ROCK Inhibitor) | Aladdin Scientific | Cat# Y129399; CAS# 129830-38-2 |
| KnockOut™ DMEM/F-12 | Gibco | Cat# 12660012 |
| N-2 MAX Media Supplement (100X) | R&D Systems | Cat# AR009 |
| MEM Non-Essential Amino Acids Solution (100X) | Gibco | Cat# 11140050 |
| GlutaMAX™ Supplement | Gibco | Cat# 35050061 |
| Doxycycline hyclate | Sigma-Aldrich | Cat# D9891; CAS# 24390-14-5 |
| SB431542 (hydrate) | Cayman Chemical Company | Cat# 13031 |
| LDN-193189 2HCl | TargetMol Chemicals | Cat# T35348; CAS# 1435934-00-1 |
| XAV-939 | TargetMol Chemicals | Cat# T1878; CAS# 284028-89-3 |
| Poly-L-ornithine hydrobromide | Sigma-Aldrich | Cat# P3655, CAS# 27378-49-0 |
| BrainPhys™ Neuronal Media | Stem Cell Technologies | Cat# 05790 |
| B-27™ Plus Supplement (50X) | Gibco | Cat# A3582801 |
| BDNF<br>Recombinant<br>Protein,<br>PeproTech® | Gibco | Cat# 450-02 |
| NT-3<br>Recombinant<br>Protein,<br>PeproTech® | Gibco | Cat# 450-03 |
| Laminin Mouse<br>Protein, Natural | Gibco | Cat# 23017015 |
| PBS, pH 7.4 | Gibco | Cat# 10010031 |
| TrypLE™ Express<br>Enzyme (1X) | Gibco | Cat# 12604013 |
| StemPro™<br>Accutase™ Cell<br>Dissociation<br>Reagent | Gibco | Cat# A1110501 |
| CryoStor® Cell<br>Cryopreservation<br>Media | Sigma-Aldrich | Cat# C287 |
| Bovine Serum<br>Albumin | Sigma-Aldrich | Cat# A7030 |
| Sodium<br>tetraborate<br>decahydrate | AMRESCO | CAS# 1330-43-4 |
| Hydrochloric acid<br>solution, 1N | Fisher<br>Chemical | Cat# SA48-1 |
| Dimethyl<br>sulfoxide, DMSO | Sigma-Aldrich | Cat# D2650 |
| Paraformaldehyde | Sigma-Aldrich | Cat# P6148; CAS# 30525-89-4 |
| Bovine Serum<br>Albumin | Sigma-Aldrich | Cat# A9647 |
| Triton™ X-100 | Sigma-Aldrich | Cat# T9284; CAS# 9036-19-5 |
| Hoechst 33342 | Invitrogen | Cat# H1399 |
| MitoBrilliant™ 646 | Bio-Techne | Cat# 7700 |
| ProLong™ Gold<br>Antifade Mountant | Invitrogen | Cat# P36930 |
| ProLong™ Gold<br>Antifade Mountant<br>with DNA Stain<br>DAPI | Invitrogen | Cat# P36931 |
| Chloroform | Fisher<br>Chemical | Cat# C298-500; CAS# 67-66-3 |
| 2-Propanol | Fisher Chemical | Cat# A416-500; CAS# 67-63-0 |
| Ethyl Alcohol | The Warner Graham Company | CAS# 64-17-5 |
| rATP, 10 mM | Promega | Cat# P1132 |
| Seahorse XF Calibrant | Agilent | Cat# 100840-000 |
| Seahorse XF 1.0 M Glucose Solution | Agilent | Cat# 103577-100 |
| Seahorse XF 100 mM Pyruvate Solution | Agilent | Cat# 103578-100 |
| Sodium chloride | Sigma-Aldrich | Cat# 71380; CAS# 7647-14-5 |
| Potassium chloride | Fisher Chemical | Cat# P217; CAS# 7447-40-7 |
| Calcium chloride dihydrate | AMRESCO | CAS# 10035-04-8 |
| Magnesium chloride | Sigma-Aldrich | Cat# M8266; CAS# 7786-30-3 |
| Potassium phosphate monobasic | Sigma-Aldrich | Cat# P0662; CAS# 7778-77-0 |
| HEPES solution | Sigma-Aldrich | Cat# H0887; CAS# 7365-45-9 |
| Sodium hydroxide solution | Fisher Chemical | Cat# SS254; CAS# 1310-73-2 |
| Critical Commercial Assays |  |  |
| Mycoplasma PCR Detection Kit | Applied Biological Materials Inc. | Cat# G398 |
| miRNeasy Micro Kit | Qiagen | Cat# 217084 |
| RNase-Free DNase Set | Qiagen | Cat# 79254 |
| TaqMan™ Reverse Transcription Reagents | Applied Biosystems | Cat# N8080234 |
| TaqMan™ MicroRNA Reverse Transcription Kit | Applied Biosystems | Cat# 4366596 |
| TaqMan™ Universal Master Mix II, no UNG | Applied Biosystems | Cat# 4440040 |
| TaqMan™ MicroRNA Assay, hsa-let-7a | Applied Biosystems | Cat# 4427975, Assay ID# 000377 |
| TaqMan™ MicroRNA Assay, hsa-let-7c | Applied Biosystems | Cat# 4427975, Assay ID# 000379 |
| TaqMan™ MicroRNA Assay, hsa-let-7f | Applied Biosystems | Cat# 4427975, Assay ID# 000382 |
| TaqMan™ MicroRNA Control Assay, RNU48 | Applied Biosystems | Cat# 4427975, Assay ID# 001006 |
| TaqMan™ Non-coding RNA Assay, RNA5S | Applied Biosystems | Cat# 4426961, Assay ID# Hs03682751_gH |
| Lactate-Glo™ Assay | Promega | Cat# J5021 |
| CellTiter-Glo® 2.0 Cell Viability Assay | Promega | Cat# G9242 |
| CyQUANT™ Direct Cell Proliferation Assays | Invitrogen | Cat# C35011 |
| Seahorse XF Cell Mito Stress Test Kit | Agilent | Cat# 103015-100 |
| Seahorse XFe96 Extracellular Flux Assay Kit | Agilent | Cat# 103793-100 |
| Deposited data |  |  |
| Raw and analyzed bulk RNA-seq data | This paper | GEO: |
| Raw and analyzed small RNA-seq data | This paper | GEO: |
| Experimental models: Cell lines |  |  |
| Human: hESC H1 (NIH approval number NIHhESC-10-0043) | Wicell | WA01 (H1) |
| Software and algorithms |  |  |
| FastQC | Andrews, 2010 | https://www.bioinformatics.babraham.ac.uk/projects/fastqc/ |
| FLASH | Magoc and Salzberg, 2011 | https://ccb.jhu.edu/software/FLASH/ |
| Cutadapt | Martin, 2011 | https://cutadapt.readthedocs.io/en/stable/ |
| HISAT2 | Kim et al., 2015 | https://daehwankimlab.github.io/hisat2/manual/ |
| featureCounts | Liao et al., 2014 | https://subread.sourceforge.net/featureCounts.html |
| BLAST+ | Camacho et al., 2009 | https://www.ncbi.nlm.nih.gov/books/NBK279690/ |
| DESeq2 | Love et al., 2014 | https://bioconductor.org/packages/devel/bioc/vignettes/DESeq2/inst/doc/DESeq2.html |
| Samtools | Li et al., 2009 | https://samtools.sourceforge.net/ |
| GORilla | Eden et al., 2009 | http://cbl-gorilla.cs.technion.ac.il |
| Zen Blue 2.3 | Zeiss | n/a |
| Fiji | Schindelin et al., 2012 | https://fiji.sc |
| Prism | GraphPad | n/a |
| Other |  |  |
| Novaseq X Plus Platform | Illumina | n/a |
| Novaseq X Platform | Illumina | n/a |
| Bioanalyzer High Sensitivity DNA Analysis System | Agilent |  |
| CFX Connect Real-Time PCR Detection System | Bio-Rad | n/a |
| BioTek Synergy H1 Microplate Reader | BioTek | n/a |
| Seahorse XFe96 Analyzer | Agilent | n/a |

## Experimental model and subject details

All studies and procedures using stem cells followed Johns Hopkins University Institutional Stem Cell Research Oversight (ISCRO) Committee guidelines. The WA01 (H1) human embryonic stem cell line was a generous gift from Dr. Lindy E. Barrett, at the Broad Institute of MIT and Harvard, and commercially obtained from WiCell Research Institute.^78^ WA01 (H1) cells were used to generate hPSCs with stably integrated iNGN2 (inducible Neurogenin-2) as previously described.^35,36,39^ Stem cell culture was carried out as previously described.^35–37^ In brief, hPSCs were cultured and passaged in mTeSR1 medium (Stem Cell Technologies) on Geltrex-coated (Gibco) plates at 37°C and 5% CO_2_. Cell lines underwent QC testing to confirm expected genotypes via STR profiling, absence of mycoplasma (Applied Biological Materials Inc.), and expression of pluripotency markers.

## Method details

### Generation of human i^3^Neurons

Human i^3^Neurons were generated essentially as previously described.^35–37,44^ In brief, H1 hPSCs with stably integrated iNGN2 (designated Day -2) were treated for three days with doxycycline (2 µg/mL, Sigma-Aldrich) to induce NGN2 expression in Induction Medium (Knockout DMEM/F-12 supplemented with N-2 MAX, MEM Non-Essential Amino Acids, GlutaMAX) on Geltrex-coated plates. SMAD inhibitors (SB431542, 10 µM, Cayman Chemical Company; LDN-193189, 100 nM, TargetMol Chemicals) and Wnt inhibitors (XAV939, 2 µM, TargetMol Chemicals) were included in the Induction Medium for assays using the Seahorse XF Analyzer to promote uniform neuronal differentiation. Subsequently, cells (designated Day 1) were replated onto Poly-L-ornithine (PLO)-coated plates and maintained in Cortical Culture Medium (BrainPhys Neuronal Media supplemented with B-27 Plus, BDNF, NT3, Laminin). Neurons were analyzed at days 2, 7, 14, and 21 of in vitro differentiation as indicated in figure legends.

### Immunocytochemistry

hPSCs and i^3^Neurons grown on 8-well glass culture slides were fixed in 4% paraformaldehyde in PBS for 20 min at room temperature. Cells were permeabilized and blocked in 0.3% Triton X-100, 5% BSA in PBS for 60 min at room temperature. Antibodies were diluted in 0.3% Triton X-100, 1% BSA in PBS, and cells were incubated in primary antibody overnight at 4°C and in secondary antibody for 2 hr at room temperature. Primary antibodies used were as follows: goat polyclonal anti-OCT3/4 (1:125, R&D Systems); mouse monoclonal anti-SOX2 (1:100, R&D Systems); mouse monoclonal anti-TRA-1-60 (1:200, Santa Cruz Biotechnology); rabbit polyclonal anti-MAP2 (1:500, Sigma-Aldrich); mouse monoclonal anti-Tubulin β 3 (1:500, BioLegend). Secondary antibodies used were as follows: Donkey anti-Goat IgG (H+L) Cross-Adsorbed Secondary Antibody, Alexa Fluor 488 (1:1000, Invitrogen); Goat anti-Mouse IgG (H+L) Highly Cross-Adsorbed Secondary Antibody, Alexa Fluor 568 (1:1000, Invitrogen); Donkey anti-Rabbit IgG (H+L) Highly Cross-Absorbed Secondary Antibody, Alexa Fluor 568 (1:500, Invitrogen). Samples were incubated in 10 mg/μL Hoechst 33342 (Invitrogen) in PBS for 10 min, for nuclear staining, and mounted with ProLong Gold Antifade (Invitrogen).

### Mitochondria Staining & Quantification

Immediately prior to fixation, cells were incubated with 200 nM MitoBrilliant 646 (Bio-Techne) in appropriate cell culture medium for 30 min at 37°C and 5% CO_2_. Cells were immediately fixed in 4% paraformaldehyde in PBS, pre-warmed to 37°C, for 20 min at room temperature. Samples proceed to permeabilization, blocking, and antibody staining as described above. For imaging, 8 confocal slices (step distance 0.24 µm) were collected, and maximum intensity projections were used for analysis. For quantification, analysis was applied to five random fields of view with a total of >40 cells quantified for each cell type. A mitochondrial mask was created using MitoBrilliant fluorescence and cell bodies were defined as areas within a 5 µm from nuclear stain (Hoechst). The fraction of mitochondrial area (%) in each cell was calculated by dividing the total area of mitochondrial mask by the cell body area. TRA-1-60 and TUJ1 were used as cell body markers in hPSCs and i^3^Neurons respectively.

### Imaging

All images were acquired using a Zeiss LSM 800 inverted laser-scanning confocal microscope and ZEN (ZEN Blue, Version 2.3).^79^ Mitochondria images were acquired using ZEN Airyscan and a 63x/1.4 NA oil objective. All other images were acquired using standard LSM settings and 10x/0.45 NA air, 25x/0.8 NA oil or 63x/1.4NA oil objectives. Imaging settings, including pinhole size, scan speed, and averaging, were kept the same between experimental groups. All presented images are maximum intensity projections generated using ZEN orthogonal projections. Representative figure generation and quantitative analysis were performed with FIJI/ImageJ software.^80^

### Quantitative reverse transcription polymerase chain reaction (RT-qPCR)

RNA was isolated using miRNeasy Micro Kit (Qiagen) with DNase digestion. 100 ng RNA from each sample was used for bulk RNA reverse transcription using TaqMan Reverse Transcription Reagents (Applied Biosystems); 10 ng RNA from each sample was used for small RNA reverse transcription using Taqman MicroRNA Reverse Transcription Kit and appropriate TaqMan MicroRNA Assay stem loop primer (Applied Biosystems). RT-qPCR was performed on a CFX Connect Real-Time PCR Detection System (Bio-Rad) using TaqMan Universal Master Mix II, no UNG and appropriate TaqMan Assay (Applied Biosystems) according to manufacturer instructions. RT-qPCR data was normalized to averaged RNU48 and RNA5S Ct values for each sample using the ΔΔCt method.

### Bulk RNA-sequencing

RNA was isolated using miRNeasy Micro Kit (Qiagen) with DNase digestion. Libraries were prepared using NEBNext Ultra II Directional RNA Library Prep Kit with poly-A isolation (New England Biolabs) and sequenced using NovaSeq X Plus Platform (Illumina). Following read processing by FastQC,^81^ FLASH,^82^ and Cutadapt,^83^ reads were then aligned to the hg38 reference genome assembly using HISAT2,^84^ annotated and quantified using featureCounts,^85^ and normalized by transcripts per million (TPM). Differential gene expression analysis of raw counts was performed by DESeq2.^86^ GO analysis was performed using GOrilla.^87^ Library preparation and Illumina sequencing were performed by the Johns Hopkins University Integrated Genomics Center (IGC). Raw and processed data were deposited in NCBI GEO and will be publicly available as of the date of publication.

### Small RNA-sequencing

RNA was isolated using miRNeasy Micro Kit (Qiagen) with DNase digestion. 500 ng of total RNA was used as input for the llumina TruSeq Small RNA Sample Kit according to manufacturer instructions. This approach uses a naturally occurring miRNA specific structure with a 5’-phosphate and a 3’-hydroxyl to ligate adapter sequences exclusively to miRNA species. After adapter ligation, libraries were prepared by reverse transcription, PCR amplification, and polyacrylamide gel electrophoresis purification. Library quality and quantity were assessed with the Agilent Bioanalyzer using the High Sensitivity Chip according to manufacturer instructions. Sequencing was performed using the Illumina Novaseq X using 150 bp by 150 bp paired end chemistry. Depth of coverage was targeted for 50 million total reads per sample. Following read processing by FastQC^81^ and Cutadapt,^83^ reads were then aligned using BLAST+^88^ and the miRBase database (Release 22.1, hg38 reference genome assembly).^89^ Reads were quantified using bash commands and normalized by counts per million (CPM). Differential gene expression analysis of raw counts was performed by DESeq2.^86^ Library preparation and Illumina sequencing were performed by the Sidney Kimmel Comprehensive Cancer Center (SKCCC) Experimental and Computational Genomics Core (ECGC). Raw and processed data were deposited in NCBI GEO and will be publicly available as of the date of publication.

### Lactate measurement

hPSCs and i^3^Neurons were grown on Geltrex and PLO- and laminin-coated 96-well plates respectively. Cell culture medium was replaced and collected 24 hr later. Medium was diluted 1:100, to fit within the linear range of the assay, and lactate measured using Lactate-Glo Assay (Promega) according to manufacturer instructions. BioTek Synergy H1 Microplate Reader (BioTek) was used to measure resulting luminescence. Lactate secretion was calculated using a standard curve and each sample normalized to cell number, determined using CyQUANT Direct Cell Proliferation Assay (Invitrogen).

### ATP measurement

hPSCs and i^3^Neurons were grown on Geltrex and PLO- and laminin-coated 96-well plates respectively. ATP concentration was measured using Cell Titer-Glo Assay (Promega) according to manufacturer instructions. BioTek Synergy H1 Microplate Reader (BioTek) was used to measure resulting luminescence. ATP concentration was calculated using a standard curve and each sample normalized to cell number, determined using CyQUANT Direct Cell Proliferation Assay (Invitrogen).

### Seahorse XF Analyzer

hPSCs and i^3^Neurons were grown on Geltrex and PLO- and laminin-coated Seahorse XFe96 cell culture microplates respectively. Approximately 1 hr before the assay, cell culture medium was replaced with Seahorse Assay Medium (Artificial Cerebrospinal Fluid (ACSF; 120 mM NaCl, 3.5 mM KCl, 1.3 mM CaCl_2_•2H_2_O, 1 mM MgCl_2_, 0.4 mM KH_2_PO_4_, 5 mM HEPES solution),^90^ 10 mM Glucose, 2 mM Pyruvate). ECAR (extracellular acidification rate, mpH/min) and OCR (oxygen consumption rate pmol/min) was measured using a Seahorse XFe96 Analyzer (Agilent), according to manufacturer instructions. Inhibitors provided in the Seahorse XF Cell Mito Stress Test Kit (Agilent) were diluted in Seahorse Assay Medium and final concentrations used were as follows: 1 μM Oligomycin, 1 μM FCCP, and 0.5 μM Rotenone/Antimycin A. Measurements were normalized to cell number, determined using CyQUANT Direct Cell Proliferation Assay (Invitrogen).

### Quantification and statistical analysis

Replicates refer to independent wells or plates, unless otherwise indicated in figure legends; all experiments include three or more independent biological replicates as noted in figure legends. Results are expressed as mean ±SD. Unpaired t-tests and ANOVA with post hoc tests were used to calculate statistical significance; p values < 0.05 were considered statistically significant. Statistical analyses were performed using GraphPad Prism 8.

