## Supplemental Information for "A human model of the neurodevelopmental metabolic shift"

**Authors and Affiliations:**

Sydney N. Pettit[1,2], Ariella T. Kornfeld[1,2], Celeste Morales[1,2] , Mollie K. Meffert[1,2,*]

[1] Solomon H. Snyder Department of Neuroscience, Johns Hopkins University School of Medicine, Baltimore, Maryland 21205, USA

[2] Department of Molecular Biology and Genetics, Johns Hopkins University School of Medicine, Baltimore, Maryland 21205, USA

**Supplemental Information**

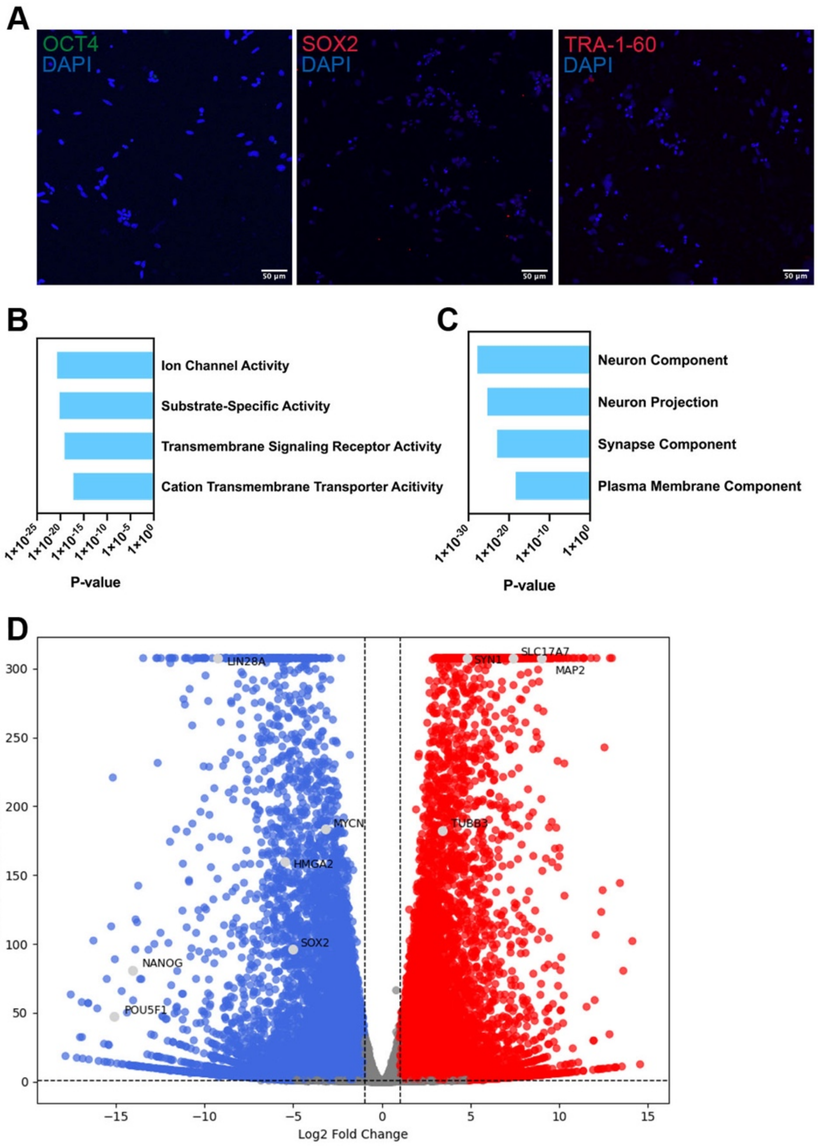


**SUPPLEMENTAL FIGURE 1: Pluripotency genes are downregulated in i$^3$Neurons.**

A) Representative ICC images showing Day 7 i$^3$Neurons negative for pluripotent markers OCT4, SOX2, and TRA-1-60.

B) Gene ontology (GO) analysis of bulk RNA-seq data showing genes up-regulated during i$^3$Neuron differentiation (Day 21); molecular functions are ranked by p-value.

C) Gene ontology (GO) analysis of bulk RNA-seq data showing genes up-regulated during i$^3$Neuron differentiation (Day 21); cellular components are ranked by p-value.

D) Bulk RNA-seq data, following analysis with DESeq2, showing up- (red) and down-regulated (blue) neuronal and pluripotency genes in Day 21 i$^3$Neurons. Highlighted genes are labeled and shown in light gray.

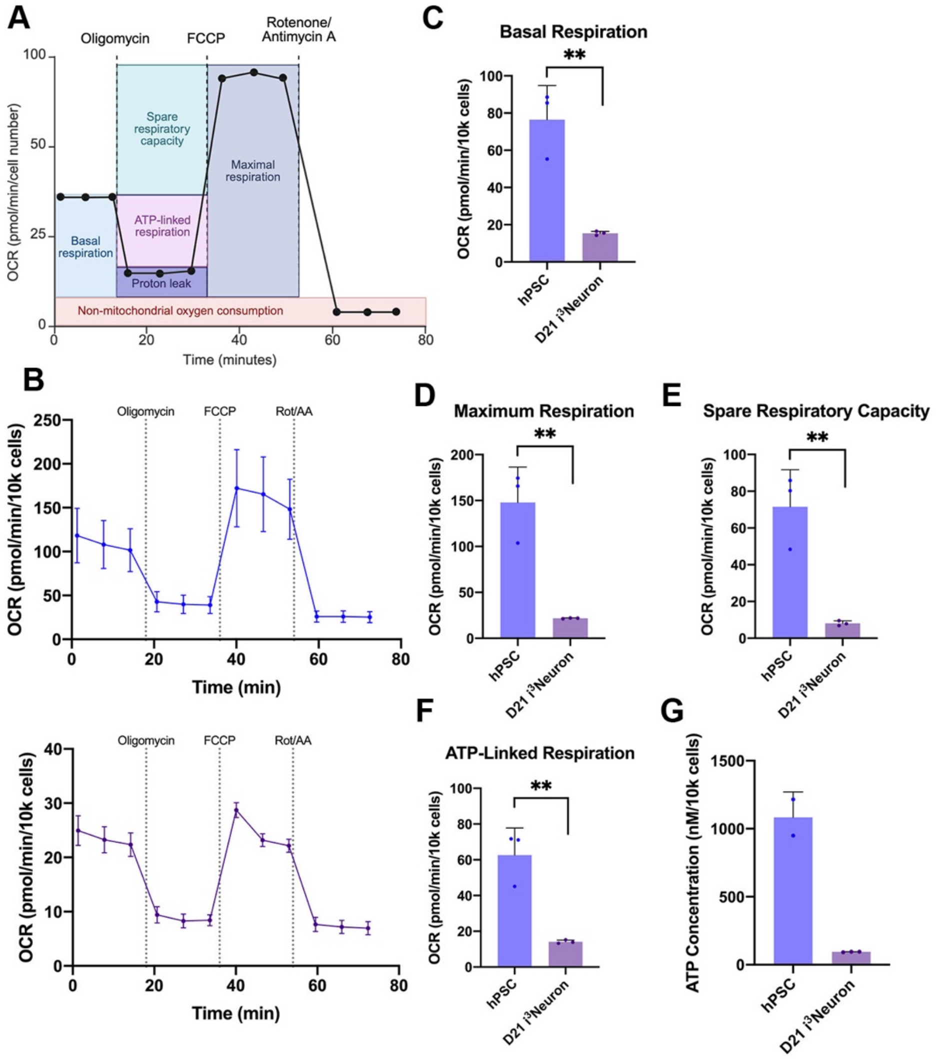


**SUPPLEMENTAL FIGURE 2: Day 21 i$^3$Neurons exhibit decreased mitochondrial respiration compared to hPSCs.**

A) Schematic depicting the Seahorse XF Analyzer Mitochondrial Stress Test which measures oxygen consumption rate (OCR, pmol/min) following injections of oligomycin, FCCP, and rotenone/antimycin A (Rot/AA) injection.

B–F) Seahorse XF Analyzer Mitochondrial Stress Test data demonstrating

B) oxygen consumption rate (OCR, pmol/min) following drug injections in hPSCs (*top*) and Day 21 i$^3$Neurons (*bottom*);

C) basal respiration (pmol/min);

D) maximum respiration (pmol/min);

E) spare respiratory capacity (pmol/min);

F) ATP-linked respiration (pmol/min)

in hPSCs and Day 21 i$^3$Neurons, normalized by cell number. Each dot represents one experimental replicate, with 30–60 biological replicates per experiment [Unpaired t-test].

G) Cellular ATP concentration (nM), measured using a Promega Cell Titer-Glo assay and normalized by cell number, in hPSCs and Day 21 $i^3$Neurons. Each dot represents one experimental replicate, with 25–30 biological replicates per experiment.
Error bars represent mean ±SD, **p< 0.01.

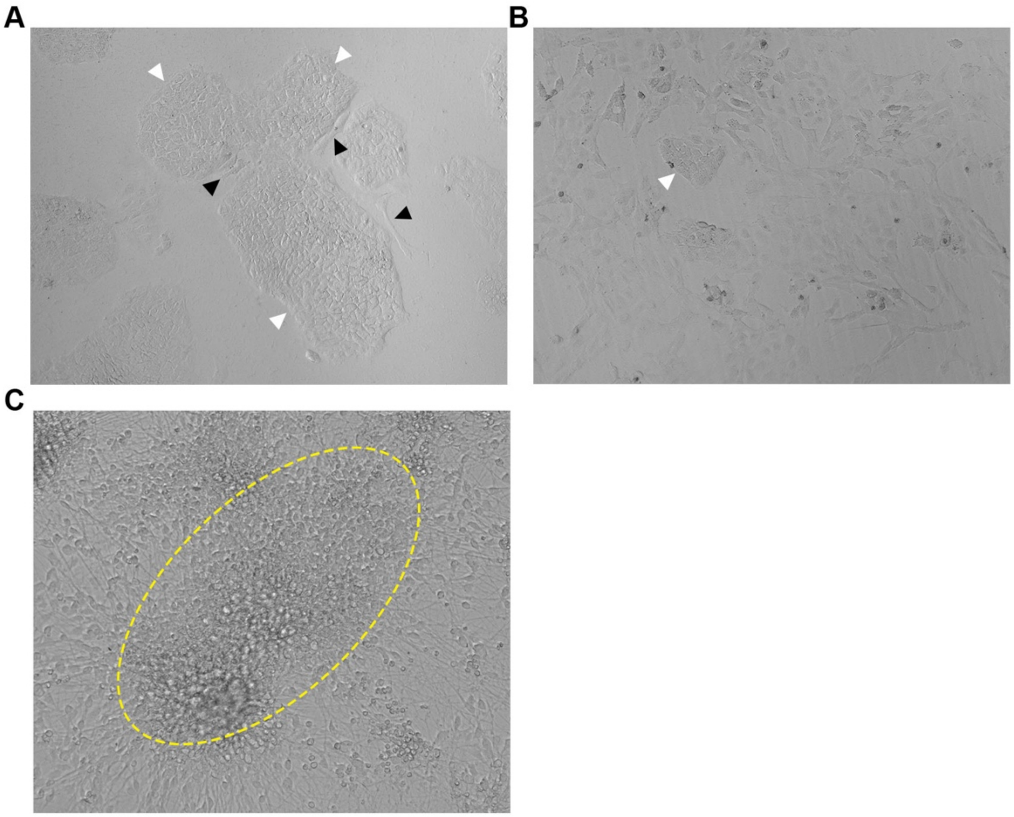


**SUPPLEMENTAL FIGURE 3: Potential problems during hPSC culture and i$^3$Neuron differentiation.**

A) Representative image of healthy hPSC colonies (white arrows) with minor spontaneous differentiation (black arrows).

B) Representative image of hPSCs with major spontaneous differentiation. White arrow indicates an undifferentiated hPSC colony.

C) Representative image of a persistent population of undifferentiated cells in Day 21 i$^3$Neuron culture (clustered cells, outlined).
